# Alignment-free identification of COI DNA barcode data with the Python package Alfie

**DOI:** 10.1101/2020.06.29.177634

**Authors:** Cameron M. Nugent, Sarah J. Adamowicz

## Abstract

Characterization of biodiversity from environmental DNA samples and bulk metabarcoding data is hampered by off-target sequences that can confound conclusions about a taxonomic group of interest. Existing methods for isolation of target sequences rely on alignment to existing reference barcodes, but this can bias results against novel genetic variants. Effectively parsing targeted DNA barcode data from off-target noise improves the quality of biodiversity estimates and biological conclusions by limiting subsequent analyses to a relevant subset of available data. Here, we present Alfie, a Python package for the alignment-free classification of cytochrome c oxidase subunit I (COI) DNA barcode sequences to taxonomic kingdoms. The package determines *k*-mer frequencies of DNA sequences, and the frequencies serve as input for a neural network classifier that was trained and tested using ~58,000 publicly available COI sequences. The classifier was designed and optimized through a series of tests that allowed for the optimal set of DNA *k*-mer features and optimal machine learning algorithm to be selected. The neural network classifier rapidly assigns COI sequences to kingdoms with greater than 99% accuracy and is shown to generalize effectively and make accurate predictions about data from previously unseen taxonomic classes. The package contains an application programming interface that allows the Alfie package’s functionality to be extended to different DNA sequence classification tasks to suit a user’s need, including classification of different genes and barcodes, and classification to different taxonomic levels. Alfie is free and publicly available through GitHub (https://github.com/CNuge/alfie) and the Python package index (https://pypi.org/project/alfie/).

## Introduction

Biodiversity is declining across the globe. Millions of species face the threat of extinction, and ecosystems are being irreversibly altered due to loss of biomass and changes in species composition (Barnosky *et al.* 2011; Ceballos *et al.* 2015). To maintain the health of ecosystems and curb biodiversity loss, informed conservation and management practices are required. Achievement of conservation goals is limited by a lack of fundamental information about species composition for many of the world’s ecosystems. It is therefore imperative that technological solutions are developed to enable the accurate and efficient characterization of the world’s biodiversity, so that existing species can be catalogued, and informed conservation strategies can be developed to protect the planet’s ecosystems.

The field of DNA barcoding offers a technological solution to the problem of taxonomically classifying organismal specimens (Hebert *et al.* 2003). Instead of relying on laborious and error-prone phenotypic classifications, sequence diversity within standardized gene regions is used to enable both specimen identification and species discovery (Hebert *et al.* 2003; Ratnasingham & Hebert 2007; Hubert & Hanner 2015). The field has advanced from the barcoding of single specimens to the bulk analysis of samples, known as metabarcoding (Hajibabaei *et al.* 2011, 2016; Taberlet *et al.* 2012; Cristescu 2014), as well as multi-marker (Stefanni *et al.* 2018) and metagenomics approaches (Cuvelier *et al*. 2010). These methods have been applied in environmental biomonitoring, where multiple species are identified at once through the collection of environmental DNA (eDNA) (Taberlet *et al.* 2012). Despite the widespread adoption of these techniques, a fundamental problem persists: the accurate and repeatable characterization of biodiversity from eDNA and bulk-sample metabarcoding data is difficult, and conclusions drawn from analyses are strongly affected by methodological decisions (Clare *et al.* 2016; Braukmann *et al.* 2019).

Environmental biomonitoring often aims to answer ecological questions through the targeted examination of a taxonomic group of interest. DNA barcodes from a group of focus are targeted using group-specific PCR primers for one or more selected marker genes in the PCR amplification step that precedes high-throughput sequencing (Braukmann *et al.* 2019; Wilson *et al.* 2019). Some commonly used primers are overly general, which results in the amplification of non-target barcodes, introducing noise into data and confounding efforts to characterize true species composition for targeted taxonomic groups (Brandon-Mong *et al.* 2015; Zinger *et al.* 2019). Additionally, intra-group PCR bias can further confound the characterization of biodiversity. The over representation of certain taxa within the target group can result in other taxa being overlooked due to poorer amplification and sequencing coverage (Elbrecht & Leese 2015).

Shotgun sequencing of eDNA overcomes the primer issues of eDNA metabarcoding but also produces substantial sequencing noise and sequences from non-standardized genomic regions (Stat *et al.* 2017; Wilson *et al.* 2019). A trade off therefore exists; shotgun sequencing overcomes the amplification bias associated with PCR, but the majority of shotgun sequencing outputs cannot be assigned even high-level taxonomic classifications with confidence (Stat *et al.* 2017; Singer *et al.* 2020). Despite present technical limitations, eDNA shotgun sequencing and other next-generation biomonitoring techniques are seeing increased adoption thanks to their potential to characterize biodiversity more broadly (Makiola *et al.* 2020). Within this next generation of biomonitoring methodologies, tools leveraging machine-learning algorithms and available data will be essential to overcoming the limitations associated with existing methods (Cordier *et al.* 2019).

The detection of the presence and abundance of species from a specific group is hampered by off-target barcodes that are amplified and sequenced in metabarcode analysis. The failure to parse target sequences effectively from off-target noise can result in erroneously inflated estimates of biodiversity (Bengtsson *et al.* 2011). Currently, the characterization of biodiversity via metabarcode samples is primarily dependent on the alignment of sequences against a pre-defined set of reference barcodes or comparison of sequences against taxon-specific models (Altschul *et al.* 1990; Wang *et al.* 2007; Bengtsson *et al.* 2011; Bengtsson-Palme *et al.* 2015). These processes limit comparison to previously characterized barcode sequences, potentially exhibiting bias against novel genetic variants. The methods are also computationally intensive, often requiring each novel variant to be compared to each reference entry. These methods would therefore be improved through the incorporation of an alignment-free pre-filtering step that allowed for target sequences to be rapidly and accurately isolated from the whole set of metabarcode output sequences using algorithms with lower computational complexity (Zielezinski *et al.* 2017). This would reduce the number of spurious barcodes and improve inflated biodiversity estimates. Additionally, the speed of analyses would be improved by limiting subsequent alignment-based analyses to the isolated target sequences.

Alignment-free methods have been widely applied in biological sequence annotation and classification problems (Zielezinski *et al.* 2017). Alignment-free comparison is defined as any method of quantifying sequence similarity that does not produce an alignment; these methods are generally less computationally intensive and can be as effective as conventional alignments (Bonham-Carter *et al.* 2014; Zielezinski *et al.* 2017). To compare sequences without alignment, features must be extracted from sequences in order to characterize their structure. One common set of alignment-free features is *k*-mer counts, where the number of occurrences of fixed length DNA words of length *k* are quantified (Crusoe *et al.* 2015). These features can be used as inputs for machine learning models trained to predict classifications such as the taxonomic designation associated with sequences (Solis-Reyes *et al.* 2018). Machine learning models that operate on *k*-mer input features have previously been applied in DNA barcode sequence classification and other predictive tasks (Kuksa & Pavlovic 2009; Langenkämper *et al.* 2014; Ainsworth *et al.* 2016; Cordier *et al.* 2017). The application of these tools is often limited to specific taxonomic classification tasks (Kuksa & Pavlovic 2009), or they rely on user-provided sets of sequence data for model training (Langenkämper *et al.* 2014).

The goals of this study were to: (1) develop a high-level alignment-free taxonomic classification tool for metabarcoding and environmental DNA marker gene data. This tool was initially designed for the kingdom-level classification of barcode sequences from the most common animal barcode, a region of the mitochondrial cytochrome c oxidase subunit I (COI) gene. (2) To achieve this, we explore different feature sets (*k*-mer sizes) and machine learning algorithms to determine the optimal machine learning architecture for alignment-free barcode classification. (3) To make the tool accessible to other researchers, we develop a Python package and command line interface to allow the alignment-free classifier to be easily deployed in future research applications. (4) Within the Python package, we also develop an application programming interface (API) to facilitate the construction of customized alignment-free classifiers for any barcode, gene, or taxonomic group of interest. Addressing these goals led to the creation of the Python package Alfie, which contains a kingdom-level alignment-free DNA barcode classifier, as well as an API to aid users in custom alignment-free classifier construction. Alfie is free and publicly available through GitHub (https://github.com/CNuge/alfie) and the Python package index (https://pypi.org/project/alfie/).

## Methods

### Data acquisition

The Barcode Of Life Data system (BOLD) (Ratnasingham & Hebert 2007) was queried to obtain all publicly available sequences for the DNA barcode: cytochrome c oxidase subunit I (COI) (https://github.com/CNuge/data-alfie). Sequences were filtered to ensure a minimum length of 300 base pairs (bp). The five kingdom-level classifications used by the BOLD database (Animal, Bacteria and Archaea, Fungi, Plant, Protist) were maintained and utilized as the labels in subsequent classifier development. As a result of BOLD’s mandate to catalogue animal biodiversity, the database displays a significant sampling bias towards the animal kingdom. To ensure that models could be trained effectively and not be biased towards animal classification, down sampling of the animal data was performed to ensure more even representation of sequences among kingdoms. Stratified sampling of animal sequences was performed to obtain a representative subsample of 0.2% of the total set of sequences available (sequences were sampled proportionally on the taxonomic level: class; a sample size of 0.2% was chosen as this yielded a set of animal sequences roughly equal to the kingdom with the second highest number of available COI barcodes, plants) (Table 1). To train models robust to variable data quality and barcode sequence coverage, each individual barcode sequence was randomly subsampled, with a 200-600 base pair subsection of the complete barcode being retained at random and subsequently utilized in model training and testing.

**Table 1.** The numbers of COI barcode sequences obtained from BOLD for each kingdom and the number of sequences retained within different data sets used in development of the Alfie package. The raw barcode counts represent the complete set of publicly available sequences for the given kingdom. The ‘Barcodes utilized’ column is the total number of sequences used in the analysis for the given kingdoms after filtering based on minimum sequence length and down sampling to decrease imbalanced representation of the different kingdoms. The breakdown of these sequences between the train, test, and validation data sets is also shown.

| Kingdom | Raw barcode count | Barcodes utilized | Train data set size | Test data set size | Validation data set size (see Table 2) |
| --- | --- | --- | --- | --- | --- |
| Animal | 1,137,552 | 23,493 | 18,189 | 4,547 | 757 |
| Bacteria and Archaea | 5,565 | 5,547 | 4,380 | 1,095 | 72 |
| Fungi | 1,407 | 1,368 | 1,038 | 260 | 70 |
| Plant | 22,638 | 22,599 | 18,017 | 4,505 | 77 |
| Protist | 5,029 | 5,026 | 4,014 | 1,003 | 9 |
| Total | 1,172,191 | 58,033 | 45,638 | 11,410 | 985 |

Prior to splitting the data into a train and test set, a validation set was created to provide a stringent test of the final models’ ability to make external predictions. From each kingdom, a complete taxonomic class was withheld to create the validation set and simulate rare or previously unseen sequences. The class withheld from each kingdom was chosen manually, with selection being based on the distribution of barcodes across the taxonomic classes of the given kingdom. Barcode distribution was variable across kingdoms, so no suitable rule-based selection method was found; classes with intermediate levels of representation within their kingdom were selected. Classes with intermediate representation levels were chosen to provide good sample sizes for subsequent classification tests without grossly detracting from the size of available training data. For the protist kingdom, two classes were selected for inclusion in the validation set due to small intra-class barcode counts. The composition of the final validation set is described in Table 2. After the validation set was withheld, the remaining data were split into a train and test (stratified split on level: kingdom), with 80% of data comprising the training set, and the other 20% being withheld as the test set (Table 2; Supplementary File S1).

**Table 2.** The taxonomic breakdown of the validation data set. For each kingdom, a taxonomic class with a near average number of sequences in the kingdom’s whole data set was chosen for exclusion from the training set and inclusion in the validation data set. The names of the taxonomic classes and the numbers of barcode sequences withheld from training and testing for subsequent validation are shown.

| Kingdom | Withheld class | Sequence count |
| --- | --- | --- |
| Animal | Diplopoda | 757 |
| Bacteria and Archaea | Flavobacteria | 72 |
| Fungi | Leotiomycetes | 70 |
| Plant | Liliopsida | 77 |
| Protist | Heterotrichea and Colpodea | 9 |

### Feature set evaluation – *k*-mer size

Following the train-test split, different sets of alignment-free features were generated, and the accuracy of kingdom-level classifications by the resulting models were tested. For barcode sequences in the training set, *k*-mer frequencies were generated for values of *k* from 1 to 6. *K*-mer frequencies (count of a given *k*-mer divided by the total number of *k*-mers counted in a given barcode) were used as model inputs, so as to standardize the scale of input values and also ensure the models were robust to inputs of different lengths. For each *k*-mer feature set, deep neural networks with five hidden neuron layers were trained and evaluated through 5-fold cross validation (neural networks implemented using the package Tensorflow Version 2.1.0, Abadi *et al.* 2016). The choice of deep neural network-based classifiers with five hidden neuron layers was based on exploratory data analysis and preliminary model construction that showed this architecture to produce effective classifiers. The number of neurons in the hidden layers of the neural network were adjusted according to the size of the input feature set (Table 3). The 5-fold loss and accuracy metrics for the neural networks with different *k*-mer inputs were compared via a one-factor analysis of variance (ANOVA) to determine if there were significant differences in classification accuracy for different feature sets (*k*-mer sizes) and to select an optimal value of *k* for further model testing.

**Table 3.** The architectures of the neural networks tested in conjunction with the different *k*-mer feature sets. For each *k*-mer feature set and corresponding neural network, the average loss and accuracy scores from 5-fold cross validation on the training data are presented. Each neural network was comprised of a dense input layer (neuron number = number of unique *k*-mers, or 4^*k*^), five hidden layers of neurons (neuron counts for each layer given in table), and a dense output layer (neuron size equal to number of classes). The input and hidden layers utilized a rectified linear unit (relu) activation function (Agarap 2018), and the hidden layers had dropout rates of 0.3. The final output layer utilized a softmax activation function, and the models were trained using an Adam optimizer (Kingma & Ba 2014), minimizing sparse categorical cross entropy.

| ***K*-mer size** | **NN hidden layers sizes** | **Average accuracy** | **Average loss** |
| --- | --- | --- | --- |
| **1** | [4, 64, 128, 32, 16] | 0.684 | 0.899 |
| **2** | [16, 64, 128, 64, 16] | 0.935 | 0.216 |
| **3** | [64, 128, 64, 32, 16] | 0.993 | 0.038 |
| **4** | [256, 128, 64, 32, 16] | 0.994 | 0.033 |
| **5** | [1024, 512, 256, 64, 16] | 0.995 | 0.047 |
| **6** | [2080, 1040, 520, 260, 130] | 0.997 | 0.023 |

### Algorithm evaluation

After selection of the optimal *k*-mer size, a series of different machine learning models were fit using the training set and optimized through a grid search of hyperparameters. Five classification algorithms were utilized: *k* nearest neighbour (KNN), support vector machine (SVM), random forest (RF), extreme gradient boosting (XGB), and deep neural network (DNN). All models were deployed using the Python programming language (Version 3.7.4). The KNN, SVM, and RF models were implemented using the package scikit-learn (Version 0.21.3, Pedregosa *et al.* 2011), the XGB model was implemented using the package XGBoost (Version 0.90, Chen & Guestrin 2016), and the DNN was implemented using the package Tensorflow (Version 2.1.0, Abadi *et al.* 2016). In order to select optimal hyperparameters and optimize performance, for each algorithm a grid search was performed using scikit-learn’s GridSearchCV function to train a series of models on the training data set using 5-fold cross validation (Supplementary File S2). Optimal hyperparameters were selected based on the highest classification accuracy. For the DNN, a custom grid search script was used, with 5-fold cross validation and several potential values for each of the models’ respective hyperparameters (Supplementary File S3).

Following the selection of optimal hyperparameter sets through the grid searches, a final version of each model was trained using the optimal set of hyperparameters and the complete training data set. Final trained models were then used to make predictions for the previously withheld test and validation sets (Table 1; Table 2). Predicted classifications were compared to true values to determine the model with the highest classification accuracy. A single optimal alignment-free kingdom-level classifier was selected for inclusion in the Alfie package based on the accuracy of predictions made on the test and validation data. Several secondary classifier characteristics were also considered to ensure model reusability. Specifically, the file size of the trained models and the time required to make predictions were quantified to ensure that the package’s memory and time requirements were not prohibitive. The Alfie package was then constructed to allow for the model to be reused in external analyses.

## Results and Discussion

### *K*-mer size

The cross-validation accuracy scores for the different neural networks and corresponding *k*-mer feature sets were compared to determine an optimal *k*-mer feature size. The results showed that the accuracy of models improved with the *k*-mer feature size, with diminishing improvements beyond *k* = 3 (Table 3; Figure 1). A one-factor ANOVA revealed the differences to be significant (p < 2e-16, F statistic = 318.3, DF_1,2_ = 5, 24), and a subsequent Tukey’s HSD test showed the accuracy of both *k* = 1 and *k* = 2 to differ significantly from all larger values of *k* but no significant differences in the performance of pairwise comparisons between *k* 3-6. A final *k* value of 4 was selected for subsequent tests, due to the insignificant differences between the values of *k* = 3 to *k* = 6 and the conservative choice to select a *k*-mer size one larger than the apparent minimal effective feature set.

**Figure 1.**
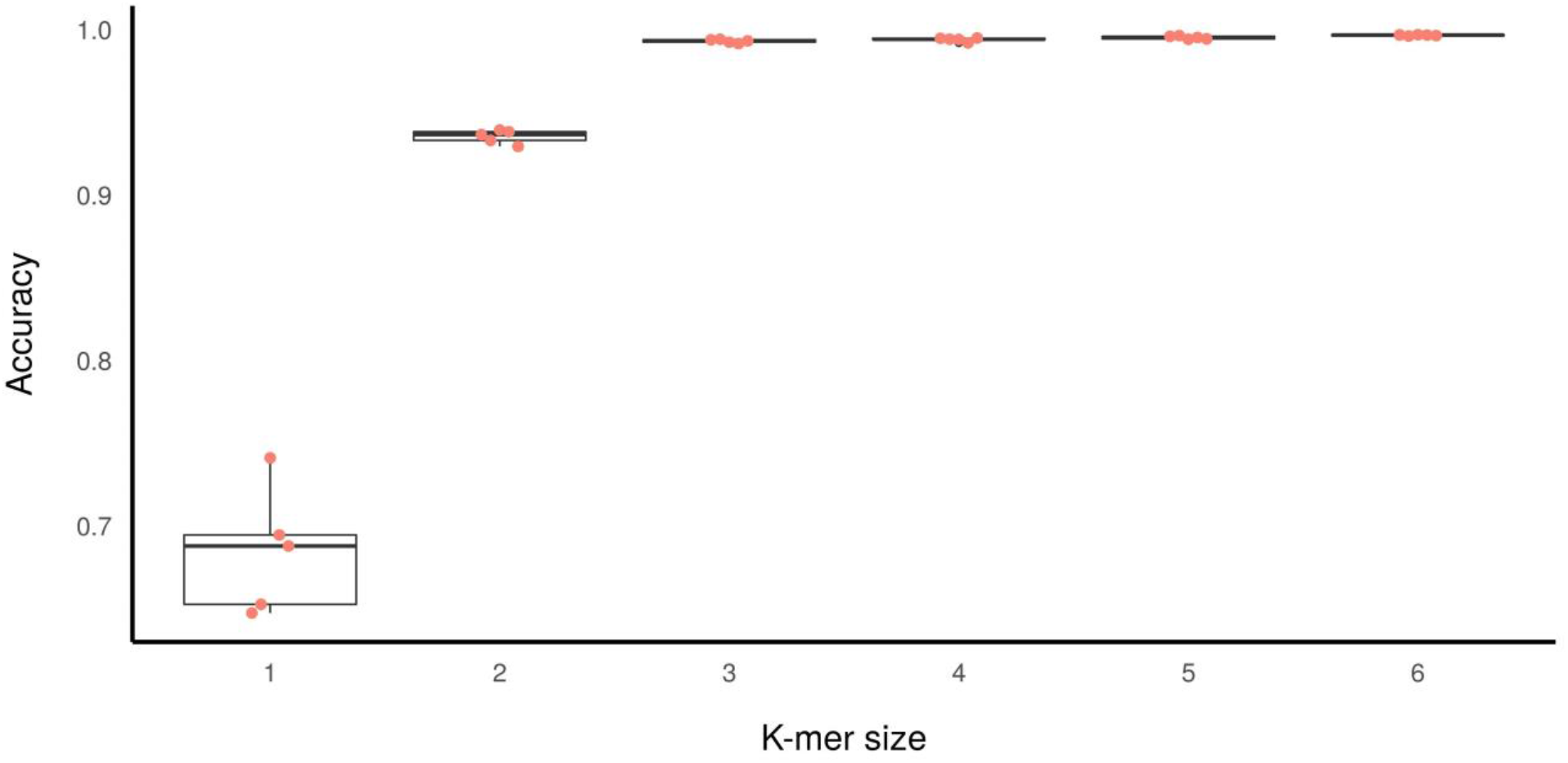
Boxplot of the 5-fold cross validation accuracy results for the training of models of different *k*-mer feature sets and corresponding neural network architectures on the training data. Each dot represents an accuracy score for one of the individual fold in the cross-validation corresponding to the given *k*-mer feature set.

### Training and validation

For each of the machine learning algorithms, a grid search was used to obtain an optimal hyperparameter set (Supplementary File S3). Final models were trained using the complete training data set and then used to make predictions for the test and validation sets (Table 1; Table 4). Performance on the test data (withheld barcodes from taxonomic groups otherwise represented in the training data) was strong for all models, with the lowest classification accuracy exceeding 98% (RF), and all other models exceeding 99.5% accuracy (Table 4). All models made less accurate kingdom-level predictions on the validation data (barcodes from taxonomic classes that were completely withheld during training) (Table 5). The accuracy was more variable across models as well. On the validation data, the accuracy score of the RF model was 0.861, and accuracy for the KNN model was 0.927, indicating poorer generalization for these methods to previously unseen data. Each of the DNN, SVM, and XGB models had accuracy >97% on the validation data, and the most accurate model was the DNN (0.976).

**Table 5.** The accuracy scores for the predictions made by the five different machine learning models (trained on 4-mer frequency features and the complete training data set). Accuracy on the test and validation data sets (Table 1) are shown.

| Algorithm | Test accuracy | Validation accuracy |
| --- | --- | --- |
| DNN | 0.996 | 0.976 |
| Support Vector Machine | 0.996 | 0.974 |
| K Nearest Neighbors | 0.997 | 0.927 |
| Random Forest | 0.983 | 0.861 |
| XGBoost | 0.998 | 0.972 |

### Final model

The DNN (operating on 4-mer input features) was selected as the final default kingdom-level classification model for the Alfie package. The DNN provided the highest accuracy on the validation data, as well as high accuracy on the test dataset. These results indicated that the model was not likely to be over fit to the training data and that it was able to generalize effectively and make predictions about data from previously unseen taxonomic classes. This generalizability of the model to rare or unseen taxa is an important feature that indicates the Alfie package can likely be used effectively in the analysis of under-studied environments where uncharacterized biodiversity is more likely to be present. The 4-mer DNN’s high accuracy on the test and validation data also indicated that the features and model can effectively capture a taxonomic signal despite no alignment being performed and variable input sequence length. The model was robust to sequences of variable lengths that spanned various subsections of the COI barcode region (variable start and stop positions in the COI barcode region, as opposed to primer-standardized sub-regions). This indicates that the alignment-free classification by Alfie is an effective method for processing DNA barcoding, metabarcoding (specific subsections of the barcode region in a given study), and potentially even applied in analysis of metagenomics data (non-standardized fragments from shotgun sequencing).

### Alignment-free model framework

The design and testing of the Alfie package presented here focuses on high-level (kingdom) classification for the most common animal barcode, COI. However, the Alfie package provides a robust framework that a user can easily apply to produce and test alignment-free classification tools for any taxonomic distinction, DNA barcode, or combination thereof (Supplementary File S4). As a kingdom-level classifier, Alfie acts as an effective data filter, allowing the barcode sequences from a kingdom of interest to be separated from the large amount of off-target noise common in metabarcode or metagenomics data. The alignment-free methods can be reapplied to further home in on taxonomic targets; for example, using publicly available data (https://github.com/CNuge/data-alfie) a binary classifier can be trained and subsequently deployed with Alfie to allow for any taxonomic group of interest to be separated from a complete set of COI metabarcode sequences. Using other publicly available data (i.e. Pruesse *et al.* 2007; Banchi *et al.* 2020), the same custom model construction and training tools in Alfie can be used to construct binary or multiclass alignment-free classification tools for other DNA barcodes or genes.

Although the Alfie package is an effective alignment-free classification framework at high taxonomic levels, traditional alignments are likely more effective for lower-level classification tasks (i.e. classification to genus or species level). The *k*-mer frequency method used by Alfie is not likely to be effective for resolving differences between closely related species with more subtle genetic differences than those seen at higher taxonomic levels. Similarly, for taxonomic groups with few representatives and no closely related outgroups, available training data may be scant, providing a limitation in training of DNNs or other machine learning models which rely on abundant training data. The integration of alignment-based and alignment-free methods for biological sequence classification has been shown to leverage the strengths of the individual approaches to yield an efficient and accurate classification method (Borozan *et al.* 2015).

A similar hybrid approach using the Alfie package for filtration of sequences and subsequent alignment of sequences for a group of interest can narrow the scope of the application of alignment methods and thereby improve both analysis speed and accuracy. The alignment-free model construction framework of Alfie can allow for multiple models to be trained with relative ease and applied in conjunction with one another to isolate barcode sequences of interest from large and messy inputs such as metagenomics data. Models could be trained and applied to: (a) separate sequences from key mitochondrial genes from other sequences, (b) assign sequences to a barcode or gene of origin, (c) conduct kingdom-level classification for different barcode genes, and (d) conduct classification at lower taxonomic levels. All this could be accomplished using the same 4-mer frequency data and would allow for messy inputs to be filtered and categorized. Processing of metagenomics data in this manner would allow subsequent alignment effort to be more strategically targeted, improving analysis speed and accuracy.

## Conclusions

We have developed and tested the Python package Alfie, which extracts *k*-mer features and uses a neural network to make kingdom-level classifications of COI DNA barcode fragments with greater than 99% accuracy. The Alfie package can therefore be used to separate barcode data for a kingdom of interest from off-target noise, narrowing the scope of subsequent analyses to only relevant data. The model is robust to full-length barcodes and short sequence fragments and is therefore an effective classifier for use in both barcode and metabarcode analyses. The Alfie package can be incorporated into broader analyses pipelines (Elbrecht *et al*. 2018; Cordier *et al.* 2019) and paired with tools that conduct quality control (Callahan *et al.* 2016; Nugent *et al.* 2020) and taxonomic annotation (Altschul *et al.* 1990; Wang *et al.* 2007) to characterize biodiversity from large and complex data sets. The default model of Alfie is limited to kingdom-level classification for the most common animal barcode, COI. Researchers may expand upon this narrow scope to fit custom research needs by using the training module of Alfie. This allows Alfie to be applied in different taxonomic classification tasks or for the classification of data from different DNA barcodes (where labelled training data are available). The generalized and customized nature of the Alfie package will allow for it to adapt along with the field of biodiversity genomics. As metagenomics becomes more prevalent, the Alfie package can be expanded with additional default models for tasks such as the isolation of mitochondrial DNA or sequences from specific mitochondrial genes from large, messy shotgun sequencing datasets.

## Supporting information

Alfie_Supplementary

## Supplementary Files

**Supplementary File S1** – Training, test, and validation data sets used in model training and analysis

**Supplementary File S2** – Python script for custom grid search of hyperparameters for optimization of the neural network.

**Supplementary File S3** – The parameters utilized in the grid search for each of the five machine learning algorithms tested in the design of the Alfie package.

**Supplementary File S4** – Jupyter notebook with tutorial demonstrating how to apply the Alfie classifier in the Python programming language, and how to train custom alignment-free classifiers using the Alfie training module.

## Acknowledgements

Thank you to Tyler A. Elliott for assisting in the acquisition of data from the BOLD database. Thank you to Christopher A. Hempel for helpful discussions during the initial conceptualization and design of the Alfie package. Thank you to Christopher A. Hempel, Rami Baghdan, and Nora Samhadaneh for feedback on the initial draft of the manuscript.

## Funding

Funding for this research was obtained from a grant in Bioinformatics and Computational Biology from Genome Canada through Ontario Genomics. Also, funding from the Ontario Ministry of Economic Development, Job Creation and Trade. Funders played no role in study design or decision to publish. This research was enabled in part by resources provided by Compute Canada (www.computecanada.ca).

## Competing Interests

The authors have declared that no competing interests exist.

## Author Contributions

The study was conceived and designed by CMN and SJA. Development of the Alfie package was performed by CMN. The initial draft of the manuscript was written by CMN. CMN and SJA contributed to the editing of the manuscript.

## Notes

https://github.com/CNuge/alfie

https://github.com/CNuge/data-alfie

### Competing Interest Statement

The authors have declared no competing interest.

