## Supplementary material for "Alignment-free identification of COI DNA barcode data with the Python package Alfie": Alfie_Supplementary: S2-DNN-parameter_grid_search.py

```
"""
currently only a cross validation here,
run a grid search and see if we can squeak a lil more performance out of the model.

"""
import numpy as np
import pandas as pd

import tensorflow as tf
from sklearn.model_selection import KFold, ParameterGrid


def dnn_class_model(hidden_sizes = [100], dropout = 0.2,
						in_shape = 4160, n_classes = 5):
	"""
	builds a simple deep neural network using the keras wrapper.
	hidden_sizes - neuron sizes for the hidden layers
				n_hidden is implict param - equal to the length of hidden layers list
	dropout - dropout applied after each hidden layer, for no dropout pass 0 
	in_shape - the number of predictors this is for 1d inputs
	n_classes - the number of output classes
	"""
	#initiate the model

	model = tf.keras.models.Sequential()
	#specify the in layer, denoting size
	model.add(tf.keras.layers.Dense(100, input_shape=(in_shape,) , activation = 'relu'))

	n_hidden = len(hidden_sizes)
	
	for i in range(0,n_hidden):
		model.add(tf.keras.layers.Dense(hidden_sizes[i], activation = 'relu'))
		if dropout != 0:
			model.add(tf.keras.layers.Dropout(dropout))


	model.add(tf.keras.layers.Dense(n_classes, activation = 'softmax'))

	model.compile(loss = 'sparse_categorical_crossentropy', 
						optimizer = 'adam', 
						metrics = ['accuracy'] )
	
	return model	

def shuffle_unison(x, y):
	assert len(x) == len(y)
	p = np.random.permutation(len(x))
	return x[p], y[p]

if __name__ == '__main__':
	
	hidden_layers = [256,128,64,32,16]

	#load in the training data
	X_train = np.load('../../train/fouronly_single_final_kingdom_X_train.npy')
	y_train_one_hot = np.load('../../train/fouronly_single_final_kingdom_y_train.npy')

	assert not np.any(np.isnan(X_train))

	X_train.shape
	y_train_one_hot.shape

	#need non-onehot encoded inputs like with the svm
	y_train = np.argmax(y_train_one_hot, axis = 1)

	# below commented out, only needed for upsample
	#note this is a bodge to overcome the fact the data are sorted - double back and fix this!
	#X_train, y_train = shuffle_unison(X_train, y_train)

	"""
	#going for a 'funnel' style nn architecture and higher dropout based off initial tests
	initial_model = dnn_class_model(hidden_sizes = hidden_layers, 
									dropout = 0.3, 
									in_shape = 256)

	#look at the model
	initial_model.summary()

	#plot the model so you can make sure its all on the up and up
	#tf.keras.utils.plot_model(initial_model, 'simple_model.png')

	#fit the data to the model
	initial_model.fit(X_train, y_train, 
						batch_size = 2000,
						epochs = 50,
						validation_split = 0.1)

	#loss going to nan

	#note - getting a vanishing gradient after about 200,000 instances passed to the model
	# tinker with the learning rate etc. to avoid this issue

	#check the outputs on the first 10 training instances
	yht_test = initial_model.predict(X_train[:10])
	#note there is a tf.argmax, old habits die hard though
	ayht_test = np.argmax(yht_test, axis = 1)

	out = initial_model.evaluate(X_train[:10], y_train[:10])
	"""

	####################################################
	# integrate this with the cv below to conduct a full cross-validation

	#TODO - add another for loop around the CV - run the different param combos
	# and record the results to a dict
	param_results = {
	'mean_test_score':[],
	'params':[],
	}


	dnn_param_grid = {'dropout' : [0.2, 0.3],
						'hidden_layers' : [[256,128,64,32,16],
											[512,256,128,32,16]],
						'epochs' : [30, 50, 70],
						'batch_size' : [100,1000,2000],}

	
	for p in list(ParameterGrid(dnn_param_grid)):

		print("on param set:")
		print(p)

		n_split = 5
		fold = 0
		cv_results = {'loss':[],
						'accuracy':[]}

		for train_index, test_index in KFold(n_split).split(X_train):
			fold_x_train, fold_x_test = X_train[train_index],X_train[test_index]
			fold_y_train, fold_y_test = y_train[train_index],y_train[test_index]
			
			model = dnn_class_model(hidden_sizes = p['hidden_layers'], 
									dropout = p['dropout'],
									in_shape = 256)

			model.fit(fold_x_train, fold_y_train, epochs=p['epochs'],
													batch_size = p['batch_size'])
			
			result = model.evaluate(fold_x_test, fold_y_test)

			cv_results['loss'].append(result[0])
			cv_results['accuracy'].append(result[1])
			

			print(f'Fold {fold} Evaluation:\n'+\
					f'loss: {result[0]}\naccuracy:{result[1]}'	)
			fold+=1

		print("cross validation results")
		cv_out = pd.DataFrame(cv_results)

		param_results['mean_test_score'].append(cv_out['accuracy'].mean())
		param_results['params'].append(p)


	grid_out = pd.DataFrame(param_results)

	grid_out.to_csv('grid_reports/4mer_dnn_grid_search_model_accuracy.tsv', index = False, 
						sep = '\t')
```
