## Supplementary material for "Alignment-free identification of COI DNA barcode data with the Python package Alfie": Alfie_Supplementary: S3-grid_paramater_lists.py

```
#nn
dnn_param_grid = {
'dropout' : [0.2, 0.3],
'hidden_layers' : [[256,128,64,32,16],
					[512,256,128,32,16]],
'epochs' : [30, 50, 70]
'batch_size' : [100,1000,2000]
}


#random forest
rf_param_grid = [{
	'n_estimators' : [500,1000,1500,2000,],
	'max_depth' : [2,6,10,None,],
	'max_leaf_nodes' : [10,16,20, None,]
	}]

#knn
knn_param_grid = [{
	'n_neighbors' : [3,5,7,9,],
	'weights' : ['uniform', 'distance'],
	}]


#SVM
svm_param_grid = [{
	'C' : [.1, 1.0, 10.0, 100.0],
	'loss' : ['hinge', 'squared_hinge'],
	}]

#xgb
xgb_param_grid = [{
	'objective' : ['multi:softmax'],
	'num_class' : [4],
	'n_estimators' :  [250,500,750,1000,],#[50, 100, 250, 400,500,600,] #500 was best on first go
	'eta' : [0.1,0.05,0.01],#[0.2, 0.1, 0.05, 0.01,],
	'max_depth' : [3,6,9],#[3, 6, 9,],
	'min_child_weight': [1,3,5],#[1,3,5,],
	'gamma' : [0,0.1,0.2],#[0,0.1,0.2,],
	}]
```
