## Supplementary material for "Alignment-free identification of COI DNA barcode data with the Python package Alfie": Alfie_Supplementary: S4-alfie_demo.ipynb

````
# %% [markdown]
# The alfie package - demo


## Part 1. Evaluate sequences with alfie's pre-built kingdom level classifier

Alfie's primary function is as a kingdom-level taxonomic classifier for COI-5P barcode data. To accomplish this, alfie uses a deep neural network to analyze a set of input sequences and predict taxonomy. 


# %% [markdown]
### Read in data 

Alfie contains a series of functions for reading and writing DNA sequence data in fasta or fastq format. These functions are found in the `seqio` module and their import is demonstrated below. Alfie also contains two example files that we import below using the read_fasta and read_fastq functions.

# %% [code]
# import functions for fasta/fastq input and output 
from alfie.seqio import read_fasta, read_fastq, write_fasta, write_fastq

# %% [code]
#import path to example file
from alfie import ex_fastq_file

#read in example file
example_fastq = read_fastq(ex_fastq_file)

#check format
example_fastq[0]

# %% [code]
#repeat the above process with a fasta file
from alfie import ex_fasta_file

example_fasta = read_fasta(ex_fasta_file)

example_fasta[0]

# %% [markdown]
### Classify sequences

Once we have imported the data with the seqio module, we can use the `classify_records` function to obtain taxonomic classifications for the input sequences.   

# %% [code]
# import classification function
from alfie.classify import classify_records

seq_records, predictions = classify_records(example_fasta)


# %% [markdown]
The classification process is completed above in just one line of code. In the background the `classify_records` function is taking the input sequences, generating a set of input features (k-mer frequencies) for each sequence, and passing the feature sets through a neural network to obtain a prediction. 

The function yields two outputs, a list of sequence records and an array of classifications. The sequence records are a list of dictionaries just like the inputs. Now there is an additional 'kmer_data' entry, which contains a data structure used to generate the input features for the neural network (more information on the `KmerFeatures` class is provided in part 2.).

# %% [code]
seq_records[0]

# %% [markdown]
The predictions returned are an encoded array of kingdom classifications.
Encodings are in alphabetical order: 
```
0 == "animalia", 1 == "bacteria", 2 == "fungi", 3 == "plantae", 4 == "protista"
```

# %% [code]
predictions[:5]

# %% [markdown]
For some purposes, the names corresponding to the encoded predictions may be preferable to the numeric encodings. To obtain these, use the `decode_predictions`function to move from the array of numeric predictions to a list of names.  

# %% [code]
from alfie.classify import decode_predictions

predicted_kingdoms = decode_predictions(predictions)
predicted_kingdoms[:5]

# %% [markdown]
Working within python provides the freedom to manipulate the sequence records in various ways using this information. What you do with the classification information will depend on your research goal. You may wish to save a select category of sequences to a file, merge the classifications with the existing sequence ids, or carry the classifications through to additional analyses in python. Here we explore a few of these possibilities.


**a.** Add the predictions into the sequence dictionaries prior to subsequent manipulation.

# %% [code]
#iterate over the predictions, use enumerate to get record number
for i, p in enumerate(predicted_kingdoms):
    #call corresponding sequence record and add a kingdom entry to dictionary
    seq_records[i]['kingdom'] = p
    
#taxonomic classification now present in the sequence dict
seq_records[0]

# %% [markdown]
**b.** Use the predictions to subset out only data from a kingdom of interest.

# %% [code]
animal_sequences = []

for i, x in enumerate(predicted_kingdoms):
    if x == 'animalia':
        animal_sequences.append(seq_records[i])
        
#Note: you could avoid the transition to string classifications and 
#subset using the numeric classifications in the `predictions` array 


# %% [markdown]
Below we see that the resulting list `animal_sequences` contains only the kingdom 'animalia'. 

# %% [code]
animal_sequences[:2]

# %% [markdown]
**c.** Writing to file

Once processing and analyses are completed, we may wish to save our sequence records to a new output file. Sequences, or lists of sequences in dictionary format can be written to output files if they possess the proper set of keys ('name' and 'sequence' for fasta, 'name', 'sequence', 'strand', and 'quality'). Additional keys will be ignored when writing the output.

In this demonstration, we take the animal sequences we isolated from the input (**b.**) and write them to a new fasta file.

# %% [code]
# if you uncomment and run the following line, it will make an output file in your current working directory
#write_fasta( animal_sequences, 'animalia_example_output.fasta')

# %% [markdown]
That is all there is to deploying the alfie package as a kingdom level classifier from within Python. The kingdom level classification provides an efficient means of separating DNA sequences from a target kingdom from the large amount of off-target noise that can exist within a metabarcoding or environmental DNA data set.

Next, we explore how the functionality of alfie can be customized to allow for isolation of target sequences on finer taxonomic scales.

# %% [markdown]

## Part 2. Train and test a custom, alignment-free taxonomic classifier

In addition to using alfie as a kingdom level classifier, alfie's helper functions can be used to train a custom DNA barcode classification model. Custom model construction will allow for the general functionality of alfie (as a kingdom-level classifier) to be extended and specialized. Some common applications of this customization may be the training of a classifier for a sub-group of interest (i.e. an intra-group classifier, which is demonstrated here for the phylum annelida), or training a binary classifier to isolate barcodes from a specific taxonomic group (i.e. a classifier that says whether an input sequence is or is not a sequence from a teleost fish).

The following demonstration will show how to train a custom binary or multiclass neural network through a combination of alfie, scikit learn, and tensorflow. alfie can be used in the training and deployment of other machine learning models, not just neural neural networks (any model that has a `predict` method is valid, which includes all models from scikit learn). For a demonstration of non neural network model deployment [see this supplementary script in the example folder](https://github.com/CNuge/alfie/blob/master/example/non_neural_net_model_example.py) that repeats the model training conducted below with a support vector machine in lieu of a neural network.

Note this process is a little more involved than the default implementation of alfie. This demo assumes the reader has an understanding of the basics of data science and machine learning in Python. If you're not yet comfortable with those topics, I would recommend the book [Hands-on Machine Learning with Scikit-Learn and TensorFlow](https://github.com/ageron/handson-ml2) as a good starting point.

It is also important to note that your mileage with a design and deployment of a custom classifier will vary with the quality of the training data you use. A few thousand labelled sequences at a minimum is recommended. If you're looking to acquire COI training data, consider: [the BOLD data systems website](http://www.boldsystems.org/index.php/Login/page?destination=MAS_Management_UserConsole), or [subsetting the data used in training the original alfie neural network](https://github.com/CNuge/data-alfie). 

If you want training data for other barcodes or genes have a look at [NCBI](https://www.ncbi.nlm.nih.gov) (warning: data mining and cleaning required), or other online barcode data sources such as the [PLANiTS dataset](https://github.com/apallavicini/PLANiTS). Another good source of barcode data is [Dr. Teresita Porter's GitHub page](https://github.com/terrimporter), which contains trained RDP Classifiers (and labelled training data!) for rbcL, 18S, ITS, and other barcodes.


# %% [code]
import numpy as np
import pandas as pd

import tensorflow as tf

from sklearn.model_selection import KFold
from sklearn.metrics import accuracy_score
from sklearn.preprocessing import LabelBinarizer

# %% [code]
from alfie.kmerseq import KmerFeatures
from alfie.training import stratified_taxon_split, sample_seq, process_sequences, alfie_dnn_default

# %% [markdown]
### Loading the demo data

The demo data can be found in the [alfie GitHub repository](https://github.com/CNuge/alfie/tree/master/example). The relative import below assumes you have downloaded the alfie repository from Github and that you are working directory is: `alfie/example`.

# %% [code]
data = pd.read_csv('alfie_small_train_example.tsv', sep = '\t')

# %% [markdown]
For simplicity, this demo is conducted with 10,000 sequences from the phylum Annelida, which has only two taxonomic classes. We will train a neural network to predict the class of Annelida sequences in an alignment-free fashion. 

# %% [code]
data.head()

# %% [code]
data['class'].value_counts()

# %% [markdown]
### Conducting a train/test split

First we sequester a test set from the input data. The aflie function `stratified_taxon_split` can be used to split a dataframe in a stratified fashion based on the taxonomic data in a column. This ensures that each taxonomic group is evenly represented in the training and test data sets.

# %% [code]
train, test = stratified_taxon_split(data, class_col = 'class', test_size = 0.3, )

# %% [markdown]
We can call some summary functions on the output dataframes to verify the even split of the data.

# %% [code]
print("train data shape:",train.shape)
print(train['class'].value_counts())
print("\n")

print("test data shape:", test.shape)
print(test['class'].value_counts())


# %% [markdown]
### Encoding the predictor data

# %% [markdown]
#### kmer features

Alfie contains a custom python class called KmerFeatures, which intakes a DNA sequence and generates k-mer count and k-mer frequency data. This class is used in the background by the `classify_records` function to generate the features for neural network prediction. 

Here we will use it to take our data and generate the feature sets for model training. The class takes an id and a DNA sequence, by default it will count 4mer frequencies.

# %% [code]
#uncomment to look at the docs
#?KmerFeatures

# %% [code]
x1 = KmerFeatures(name = train['processid'][0]  , sequence = train['sequence'][0])
x1

# %% [markdown]
Upon initiation, the KmerFeatures class instance will generate a k-mer count dictionary, where the keys are all the nucleotide permutations for size 'k'. The class then iterates through the input sequence and counts the occurrences of each k-mer. Any occurrences of nucleotides other than A, T, G, and C will cause the encompassing k-mer to be ignored.

After initiation, the k-mer keys and values can be accessed like a regular python dictionary.

# %% [code]
x1.keys()[:10]

# %% [markdown]
[Note/digression: later on, we rely on the fact keys are from an alphabetically ordered dict (the default for python 3.6 or newer). Alfie sorts the dict items by key before returning them just to be safe (Python 3.6 and later are ordered dicts by default so this is redundant). This is a PSA that it is 2020 and time to update if you're on 3.5 or earlier!](https://twitter.com/raymondh/status/773978885092323328)

# %% [code]
x1.values()[:10]

# %% [code]
x1.items()[:10]

# %% [markdown]
So that the training data is not biased by the overall size of the sequences, it is best to train the models using the k-mer frequencies (count/total), which the KmerFeatures class also provides for us.

# %% [code]
x1.freq_values()[:10]

# %% [markdown]
For machine learning purposes, the KmerFeatures class also outputs the k-mer dictionary keys and value frequencies as numpy arrays.

# %% [code]
x1.labels[:5] #only showing first 5 

# %% [code]
x1.kmer_freqs[:5] #just static method version of x1.freq_values()

# %% [markdown]
By passing the optional `k` argument, we can overrule the default k-mer size of 4 and specify a custom k-mer dictionary size. 

# %% [code]
# count 5mers 
x5mer = KmerFeatures(name = train['processid'][0]  , sequence = train['sequence'][0], k = 5)

print("start of 5mer counts:")
print(x5mer.items()[:5])

#generate 5mer and 1mer frequencies
x1_mer = KmerFeatures(name = train['processid'][0]  , sequence = train['sequence'][0], k = 1)
print("1mer counts:")
print(x1_mer.items())


# %% [markdown]
#### Random subsampling

The k-mer encoding of the predictor data above utilizes whole barcode records. Often in analysis of metabarcode or eDNA research we are not dealing with complete barcode sequences, but rather short sequences (from primer-defined barcode sub-sections in the case of metabarcoding data, or undefined sub-sections in the case of metagenomics data). 

To train our classification model on data that more closely resembles real world sequences, we can use the alfie's `sample_seq` function to randomly subsample the training sequences. We specify the `min_size` and `max_size` of the subsequence and the function will randomly generate a subsample of the sequence in the defined size range.

# %% [code]
#?sample_seq

# %% [code]
#demonstrate on a single sequence - i.e. row 1 of the train

sub_seq = sample_seq(train['sequence'][0])
sub_seq

# %% [markdown]
We can also use the sample seq function upsampling of data as well. If we enter an integer value for the  `n` argument to the function then `n` random subsamples will be generated. The function therefore returns a list of strings.

# %% [code]
sub_seq = sample_seq(train['sequence'][0], n = 10)
sub_seq[:2] #different random subsections of the same input

# %% [markdown]
### Batch processing and model input generation

We can batch process sequences, taking random subsamples, generating `KmerFeatures` class instances, and extracting the k-mer frequencies from the objects using the `process_sequences` function. This function takes in a dataframe that contains the ids, sequences and labels in different columns. The function will generate a dictionary that contains four lists of: ids, sequences, k-mer frequency arrays, and labels for each sequence.

Here both the train and test data are processed with the default k-mer size (`k = 4`) and a single subsample per sequence in the input dataframe (`n = 1`).

# %% [code]
#?process_sequences

# %% [code]
train_kmer_data = process_sequences(train, label_col = 'class')
test_kmer_data = process_sequences(test, label_col = 'class')


# %% [code]
test_kmer_data.keys() #four keys, each with lists of equal size containing the data

# %% [markdown]
The output dictionaries of the process_sequences function can be easily turned into numpy arrays, which are compatible inputs for most machine learning algorithms.

# %% [code]
print("building X arrays")
X_train = np.array(train_kmer_data['data'])
X_test = np.array(test_kmer_data['data'])

# %% [markdown]
### Encoding the response data
In addition to the alignment-fre predictor data, we also need to generate our response data. We can use scikit learn's `LabelBinarizer` function to numerically encode the taxonomic information.


# %% [code]

print("encoding y arrays")
#encode the y labels
y_train_raw =  train_kmer_data['labels']
y_test_raw = test_kmer_data['labels']

tax_encoder = LabelBinarizer()

y_train = tax_encoder.fit_transform(y_train_raw)
y_test = tax_encoder.transform(y_test_raw)

print("names are encoded in alphabetical order:")
print(tax_encoder.classes_)

# note since this is a binary class the y arrays contain only 0 and 1
# if multiclass, we need to take the argmax for multilabels 
#y_train = np.argmax(y_train, axis = 1)
#y_test = np.argmax(y_train, axis = 1)

print(y_train[:10])
print(y_train[-10:])
print(y_train.shape)
print(y_test.shape)

# %% [code]
## uncomment these lines to save all the generated arrays 
#print("saving X to files")
#np.save('X_train.npy', X_train)
#np.save('X_test.npy', X_test)

#print("saving y to files")
#np.save('y_train.npy', y_train)
#np.save('y_test.npy', y_test)

##can then load with np.load()

# %% [markdown]
### Training the neural network

After the train and test data are generated, we can use the `alfie_dnn_default` function to train a generic tensorflow neural network. We just need to specify the size of the hidden neuron layers, the shape of the input, and the number of classes we are predicting. The default dropout rate of 0.2 can be changed but will suit most situations. If you want to exert more control over the network architecture, then feel free to build your own and skip down to the section `Deploying custom trained models` for integrating it with alfie.

# %% [code]
#?alfie_dnn_default

# %% [code]
annelida_params = {
    'hidden_sizes' : [16,64,128,64,16], # the number of neurons in our hidden layers 
    'dropout' : 0.2, #the dropout rate for the neural networl
    'in_shape' : 256, #default for 4mers, can determine this through the shape of the X dataframe (# columns)  
    'n_classes' : 2, #we have two annelida classes. Default is 5 (number of kingdoms).    
}

# %% [code]
annelida_classifier = alfie_dnn_default(**annelida_params)

# %% [code]
#look at the neural network architecture
annelida_classifier.summary()

# %% [markdown]
Below the classifier is trained in an extremely simple fashion, the data is passed through the model in batches of 100 for 50 epochs. We could improve this model with cross validation and hyperparameter tuning (i.e. changing the dropout or the number of hidden layers). Additionally you could change the set of input features (size of k), or take lower level control and design a neural network yourself. These refinements and optimizations are outside of the scope of the alfie tutorial, but will offer meaningful improvement in most situations. Also scikit learn and tensorflow have excellent functions and documentation if you wish to branch out beyond the `alfie_dnn_default` architecture.

# %% [code]
annelida_classifier.fit(X_train, y_train, 
                        batch_size = 100,
                        epochs=50, 
                        verbose = 0) #switch to verbose = 1 if you want to watch it train

# %% [markdown]
After training the model, we can use it to make predict classifications for our test data. Below we make these predictions, and then compare them to the known values to get an external measure of the annelid classifier's accuracy. Even with such a simplistic training approach we have produced a classifier that is >98% accurate.

# %% [code]

#make preditions
yht_dnn_vals = annelida_classifier.predict(X_test)

#get the argmax of the predictions
yht_dnn = np.argmax(yht_dnn_vals, axis = 1)

#evaluate against the true values
dnn_score = accuracy_score(y_test, yht_dnn)

print("accuracy of model on classifying the test data:")
print(dnn_score)

# %% [markdown]
We can use the alfie `decode_predictions` function to move from binary classifications back to taxonomic names, just pass the list of names in alphabetical order (here these are lazily obtained using the original encoder's class list).

# %% [code]
y_test_class_predictions = decode_predictions(yht_dnn, tax_list = tax_encoder.classes_)

y_test_class_predictions[:10]

# %% [markdown]
### Deploying custom trained models

Once you have optimized a custom model, you can save it to a file and then reuse it on novel data by employing alfie's `classify_records` function.

# %% [code]
##save and load via tensorflow

##save model to file
#annelida_classifier.save('annelida_demo_classifier')
##load model from file
#annelida_classifier = tf.keras.models.load_model('annelida_demo_classifier')


# %% [markdown]
Here we move from the test dataframe to the list of dictionary format returned by alfie's `read_fasta` format (simulating reading new data in from a fasta file and evaluating it with our custom classifier). 

# %% [code]
#new list of dictionaries
test_simulated_fasta = []

#only doing first 10 rows
for i in range(0, 10):
    x = test.iloc[i]
    #make new dictionary entry for the sequence
    new_record = {'name' : x['processid'], 
                 'sequence': x['sequence']}
    #append to the list
    test_simulated_fasta.append(new_record)

#observe the data format
test_simulated_fasta[0]

# %% [markdown]
The `classify_records` function can be used on data in this format, we just add the additional argument `dnn_model` to use the custom annelida classifying neural network as opposed to the default kingdom-level classifier. Our custom model uses a k-mer size of 4, so this parameter is not changed. For models trained on data for other k-mer sizes, the `k` argument must also be set to the correct value.

# %% [code]
test_out, test_predictions = classify_records(test_simulated_fasta, dnn_model = annelida_classifier)

# %% [markdown]
As shown in part 1, this function yields the input sequences with the kmer_data added to their records, and a list of predictions.

# %% [code]
print(test_out[0])

print("predicted values:")
print(test_predictions)

# %% [markdown]
The decode predictions function can then be used to move from numeric predictions back to class labels.

# %% [code]
decode_predictions(test_predictions, tax_list = tax_encoder.classes_)

# %% [markdown]
The custom models can be used from outside of python as well in conjunction with the alfie command line function! Simply specify the custom model with the `-m` flag and the custom classes with the `-c` flag to conduct custom file-to-file classification.

# %% [markdown]
A few closing notes on the nuances and limitations of custom model training:

- The alfie classification method is rapid, and accurate for higher taxonomic classes. It is however not a complete replacement for sequence alignment as it lacks the detail provided by sequence-to-sequence comparison. The initial purpose of alfie was to isolate sequences of interest from larger data sets and to narrow the search space for subsequent alignments. Use the package wisely!
- The number of training samples and quality of data will influence how well a custom classifier performs, the performance of custom classifiers therefore cannot be guaranteed.
- A common issue with multiclass taxonomic classification is class imbalance. For example you may have a phylum that contains 6 classes with the following sample counts:
    ```
    class_a = 2378
    class_b = 1411
    class_c = 1219
    class_d = 377
    class_e = 27
    class_f = 12 
    ```
  With so few training instances for classes e and f, the neural network you build will likely classify individuals from these sparsely represented classes more poorly. In situations like this, you will have to get creative and try a few different things out. Possible solutions to this problem could be: upsampling the low frequency classes (using the `sample_seq` function), grouping the classes with smaller sample sizes into an 'other' category or adding them to the class of their closest taxonomic relatives, or possibly a series of binary classifier. No one of these solutions will work in all situations, so the best approach will depend on both your research goal and the data you're using.
````
