## Supplementary material for "Alignment-free identification of COI DNA barcode data with the Python package Alfie": Alfie_Supplementary: data_split_and_kmer_no_up_4mer.py

```
import gc
import numpy as np
import pandas as pd

from processing_files.kmerseq import KmerFeatures
from processing_files.seqsubsetting import seq_samples

from sklearn.preprocessing import LabelBinarizer
from sklearn.model_selection import StratifiedShuffleSplit


def stratified_train_test_valid_taxon(input_data, class_col, test_size = 0.3):
	"""
	take a dataframe and conduct Stratified a train/test split based of a 
	user defined categorical column
	"""
	print(f'Conducting train/test split, split evenly by: {class_col}')

	#split off a test/valid set, 30% of the data total
	strat_index = StratifiedShuffleSplit(n_splits=1, test_size=test_size, random_state=1738)

	for train_index, test_valid_index in strat_index.split(input_data, input_data[class_col]):
		X_train, X_test = input_data.loc[train_index], input_data.loc[test_valid_index] 


	return X_train, X_test


def upsample_fragments(seq_df, label_col = 'kingdom',  kmers=[6, 3] ,**kwargs):
	"""
	take in a dataframe with dna in a column names 'clean_dna', conduct repeated
	subsampling of the sequence and generate kmer information for each subsample.

	- you can control the subsampling protocol by passing seq_samples keyword arguments
	to this function.

	returns a dict of lists: 
	keys(ids, label, data)

	ids - the processids
	label - the kingdom column by default, this can be changed as needed 
	data - the kmer array is 6mer+3mer as per previous
	"""

	#stores tuples of (processid, kingdom, kmer_freqs)
	samples = {'ids': [],
				'labels': [],
				'data': [],
				'seq': []}
	# data are split to train and test
	# now do the upsampling and generation of the output data files.
	for entry in seq_df.iterrows():
		processid =  entry[1].processid
		label = entry[1][label_col]
		seq =  entry[1].clean_dna

		sub_seqs = seq_samples(seq, **kwargs)

		for s in sub_seqs:
			k_seq = KmerFeatures(processid, s, kmers=kmers)
			samples['ids'].append(processid)
			samples['labels'].append(label)
			samples['data'].append(k_seq.kmer_freqs)
			samples['seq'].append(s)

	return samples


if __name__ == '__main__':
	

	all_data = pd.read_csv('../data/alldat_final_kingdom.tsv', sep = '\t')

	all_data.shape #(57147, 10)

	all_data['seq_len'] = all_data['clean_dna'].apply(lambda x: len(x))

	all_data = all_data[all_data['seq_len'] > 300]

	all_data.shape #(57048, 10)

	all_data.kingdom.value_counts()

	all_data = all_data.reset_index(drop=True)

	train, test = stratified_train_test_valid_taxon(all_data, 'kingdom', test_size = 0.2)

	#train = train.head(100)
	#test = test.head(100)
	print("sampling test data")
	test_samples = upsample_fragments(test, kmers=[4], n=1)
	test = []
	gc.collect()

	print("sampling training data")
	train_samples = upsample_fragments(train, kmers=[4], n=1)
	train = []
	gc.collect()

	print("building X arrays")
	X_train = np.array(train_samples['data'])
	X_test = np.array(test_samples['data'])
	
	print("saving X to files")
	np.save('../train/fouronly_single_final_kingdom_X_train.npy', X_train)
	np.save('../test/fouronly_single_final_kingdom_X_test.npy', X_test)

	X_train = []
	X_test = []
	gc.collect()

	print("encoding y arrays")
	#encode the y labels
	y_train_raw =  train_samples['labels']
	y_test_raw = test_samples['labels']

	tax_encoder = LabelBinarizer()

	y_train = tax_encoder.fit_transform(y_train_raw)
	y_test = tax_encoder.transform(y_test_raw)

	y_train.shape
	y_test.shape

	print("saving y to files")

	np.save('../train/fouronly_single_final_kingdom_y_train.npy', y_train)
	np.save('../test/fouronly_single_final_kingdom_y_test.npy', y_test)

	print("saving subset data copies")
	del train_samples['data']
	del test_samples['data']

	train_lables = pd.DataFrame(train_samples)
	test_labels = pd.DataFrame(test_samples)

	print("writing subset data to files")

	train_lables.to_csv('../train/fouronly_single_final_kingdom_labels.tsv', sep = '\t', index = False)
	test_labels.to_csv('../test/fouronly_single_final_kingdom_labels.tsv', sep = '\t', index = False)
```
